# A Big Data COVID-19 literature pattern discovery using NLP

**DOI:** 10.1101/2022.06.01.494451

**Authors:** Panayiotis Petousis, Vasilis Stylianou

## Abstract

As our collective knowledge about COVID-19 continues to grow at an exponential rate, it becomes more difficult to organize and observe emerging trends. In this work, we built an open source methodology that uses topic modeling and a pretrained BERT model to organize large corpora of COVID-19 publications into topics over time and over location. Additionally, it assesses the association of medical keywords against COVID-19 over time. These analyses are then automatically pushed into an open source web application that allows a user to obtain actionable insights from across the globe.

## 1. Introduction

Since 2020, COVID-19 pandemic has led to a total of 522 million cases, resulting in 6.27 million deaths [1]. At the pandemic onset, the knowledge about the coronavirus was limited (e.g., the cause and the transmissibility, etc.), yet it rapidly spread across different countries. The science communities across the globe therefore published new discoveries related to coronavirus at an unprecedented rate, in hope to provide scientific evidence to develop vaccines and treatments to cease the spread of the virus. Within only a single year in 2020, there were more than 30,000 of the COVID-19 preprint research published [2].

Due to the large volume of information in the science community about COVID-19 and the rapid evolution of the disease, it is difficult to extract meaningful and non-contradictory information from the tremendous pool of scientific papers in an efficient manner [3]. During this period of rapid scientific research discovery, different tools have been developed to assist researchers to extract useful information from the literature [4]. Additionally, the COVID-19 pandemic resulted in public and healthcare policies across the world to contain the disease spread and to alleviate pressure from healthcare institutions. The change of healthcare policies (e.g., focusing on novel treatments) was due to new discoveries of treatments and virus variants. While this change may reduce the risk and stress on healthcare systems, it was until two years after the pandemic that the scientific community had a better picture on which individuals are at high risk for mortality [5-7].

In this work, we present a scalable framework to analyze published COVID-19 literature. The proposed framework automatically categorizes the publications into topics of interest over time and assesses association strength between an outcome (i.e., COVID-19) and a specific query (i.e., keywords) of interest. Thus, generating quick and actionable insights that normally takes hours for a reader to extract. The framework was integrated into a COVID-19 insights dashboard, which is publicly available to review our findings^1^. We hypothesize that the proposed dashboard allows clinicians to quickly identify relevant medical information globally. The organization of this paper is as follows: section 2 describes the proposed framework for insights extraction, section 3 presents the use cases of the online dashboard which is powered by the proposed framework, and section 4 discusses the strengths and limitations of the proposed method, followed by concluding remarks.

## 2. Methods

### 2.1. Materials

The data used for this analysis was extracted from dimensions.figshare.com [8] - a search engine to retrieve COVID-19 related materials, including published peer-reviewed articles, preprints, grants, and clinical trials. In this work, a total of 981,180 papers, 33,345 grants, and 63,008 clinical trial publications were retrieved to train and evaluate the models.

### 2.2. Publication grouping into medical categories

Different categories of information were identified in each source of publications, such as risk factors, treatment, vaccine, and kidney disease [9]. To extract the information, a publication would first need to be classified into a medical category so that relevant information could be retrieved. The proposed systematic approach to classify a publication is as follows: (1) a dictionary of medical keywords was first created by medical experts; (2) abstracts of all publications were extracted; (3) standard natural language preprocessing steps were applied to each abstract using NLTK [10] to obtain a clean paragraph representation for each publication. The preprocessing steps includes stopword removal, word tokenization, case standardization, and lemmatization; (4) a keyword matching approach (using the medical expert dictionary from step (1)) was applied to each publication to classify its category, i.e., a publication would be classified as category A (e.g., risk factor) if most identified keywords in the publication belong to category A. Table 1 shows the examples of categories and their corresponding keywords. These groups of keywords are then enhanced automatically using the Medical Subject Headings (MeSH) controlled vocabulary [11].

**Table 1.**
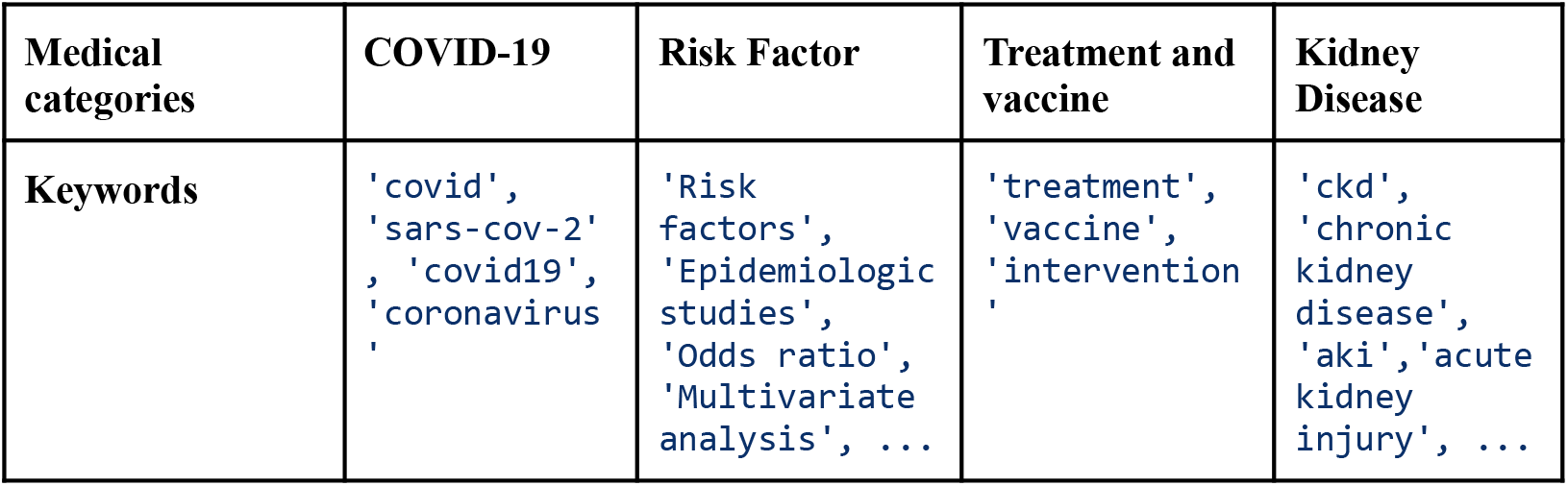
Medical categories and expert defined keywords. The complete list of keywords can be found in our public repository^2^.

### 2.3. Topic Modeling

Topic modeling is a class of unsupervised techniques that can be applied to massive unstructured documents to generate ”topics” that group similar documents. It does not require any labeling on the documents. Latent Dirichlet allocation (LDA) is a probabilistic approach of topic modeling, which is based on the assumption that all documents belong to topics. The goal of LDA is to discover the topics from documents using probabilistic modeling. The documents and their words are observed, while the topic structure, the pre-document topic distributions, and the per-document per word topic assignments belong to the hidden structure [12]. Generally, the hidden structure can be estimated using sampling-based algorithms, such as Gibbs Sampling or Variational Methods.

In this study, we applied topic modeling to learn the topics in an unsupervised manner. Initially, the text preprocessing steps described in section 2.2 were applied, then bigrams of the preprocessed texts were generated. Finally, Mallet [13], an open-sourced Java-based tool, was used to train an LDA model using the Gibbs sampling technique. A grid search approach was used to identify the optimal number of topics in each medical category, which optimized the chosen evaluation metric (the coherence score). The term-frequency and inverse term frequency (tf-idf) was used to estimate the importance of each word in the training corpus. Subsequently, we used a range of percentiles to keep the highest scoring words in our corpus before applying topic modeling. The range of the hyperparameter tuning for the tf-idf is between 0.10 and 0.90. The topic modeling algorithm was set to run for 1,000 iterations for each model fit. Each grid search was run over a range of topics automatically defined by our method based on computing an optimal number of topics from data. The range of topics had a predefined lower and upper bound of two and twelve (e.g., if the optimal number of topics was 14, then the optimal was set to 12), respectively. To compute the optimal number of topics, we used tf-idf to generate a numerical statistic of the importance of each word in our corpus. Subsequently, we used Singular Value Decomposition (SVD) with a sufficiently large number of components (e.g., 100) to reduce the dimensionality of the tf-idf matrix. Then we applied a mixture of Guassians (MoG) models on the SVD output over a range of components corresponding to the aforementioned lower and upper bound, to cluster the data and hence simulate a set of topics. Using the Bayesian Information Criterion (BIC), the MoG with the lowest BIC was used to define the optimal number of clusters (i.e., MoG with number of components that minimized BIC). Once the optimal number of components was defined, the grid search was tested with a range of ±2 topics from the optimal number. For each fitted topic model a coherence score (i.e., a score used to assess the quality of the learned topics) was computed, the model with the highest coherence score was chosen as the best model.

### 2.4. Relationship Extraction

Using the OpenNRE [14] open source model we extracted relation facts for each medical category with respect to COVID-19. An association strength estimate of each keyword in each medical category was assessed with COVID-19 using the ”wiki80 bert softmax” model. This model was pre-trained on the ”wiki80” dataset with a BERT encoder. The association strength is represented as a scalar value in the range of 0 *−* 1.‘ All our code is publicly available^3^.

### 2.5. Visualization

To visualize and share our findings, we built an open source web application using Dash [15], which is an open source framework developed by Plotly inc. The web application consists of four main pages: the “home” page, the “topic modeling” page, the “neural relationship extraction (NRE)” page, and the “about us” page. The “home” page describes the project. The “topic modeling” page depicts our analysis of topics over each dataset and each medical category across time. The “NRE” page depicts the associations of medical categories’ keywords and COVID-19 over time. The “about us” page shows information about the developers of this project.

## 3. Results and Discussions

Using an open source web application allows users to generate results on each dataset for each medical category dynamically. The web application consists of four datasets: a clinical trials dataset, a grants dataset, the top 5% most impactful papers according to altmetric, and the top 5% to 10% most impactful papers according to the altmetric score of each paper. Figure 1 depicts the distribution of topic models over the years of 2020 and 2021. The time period, location, medical category, and dataset are parameters defined by the user for the visualization of the distribution of topics and their set of keywords in the table shown in Figure 1. Some keywords are social vulnerability index (svi), medical debt. Figure 2 shows the topic distribution over time. For the top 5% most impactful papers, Topic 1 was the most dominant topic (i.e., papers belonging to topic 1 were mostly published). NRE was used to estimate the association strength of keywords over documents in each corpus. Figure 3 depicts the averaged strength of associations of each keyword across all documents per day. Figure 4 depicts the averaged strength of association over time. Each time point on Figure 4 represents averaged strength of association across multiple papers. Figure 5 depicts the averaged strength of association of keywords grouped into high level groups of keywords according to MesH, over time. This depicts how allocation of research efforts was outlined overtime. Our proposed framework allows the analysis of multiple corpora over time. Topics are generated in an unsupervised manner that allows a user to identify the predominant trend (i.e., topic) in a corpus, of each country or worldwide over time. Additionally, using the relationship extraction module a user can assess the association of certain keywords over time. For this analysis we only used the abstracts of published documents to handle the unprecedented volume of published articles. The full text can also be used for analysis. This would require an exponential increase in the computational power needed to analyze the corpus and visualize the results. The web application was deployed using the free service offered by Heroku [16], an online hosting service. Free hosting only provides limited resources, as such we only used the top 10% most impactful papers from our data source. The web application is designed to handle the full corpus of papers as well as additional literature such as the open source COVID-19 literature from semantic scholar [17], which consists of more than a million published papers.

**Figure 1.**
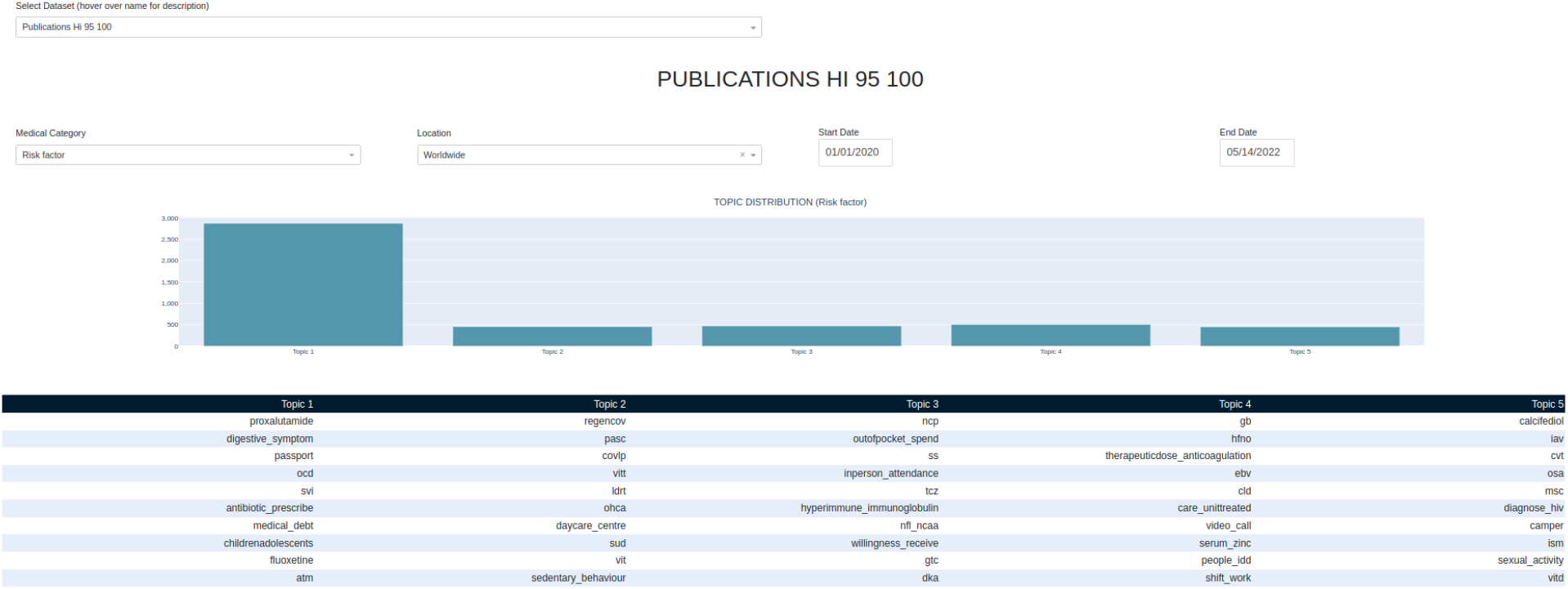
Topics distribution counts in corpus. Topic 1 is the dominant topic. Table shows the group of keywords per topic. Dates allow the selection of a time period of published articles. The location dropdown menu allows the selection of articles published in a specific country or Worldwide.

**Figure 2.**
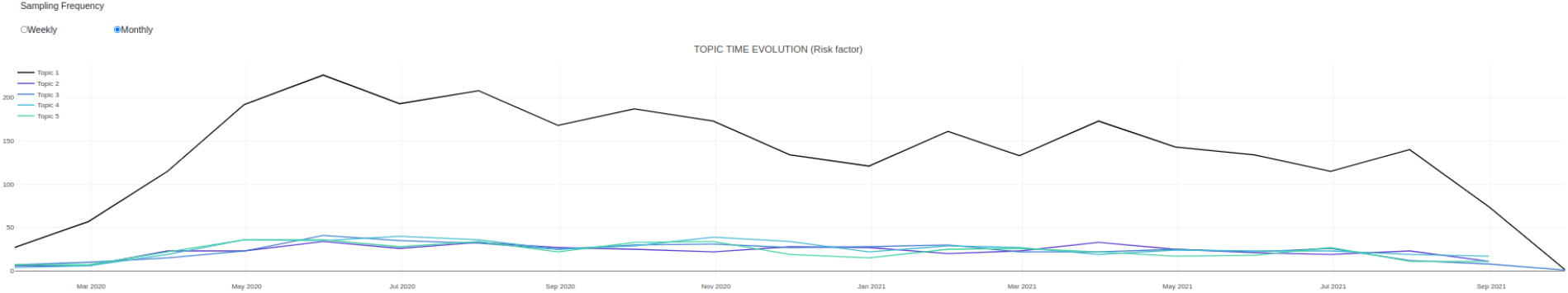
Topics distribution of published articles over time. Topic 1 is the dominant topic over time.

**Figure 3.**
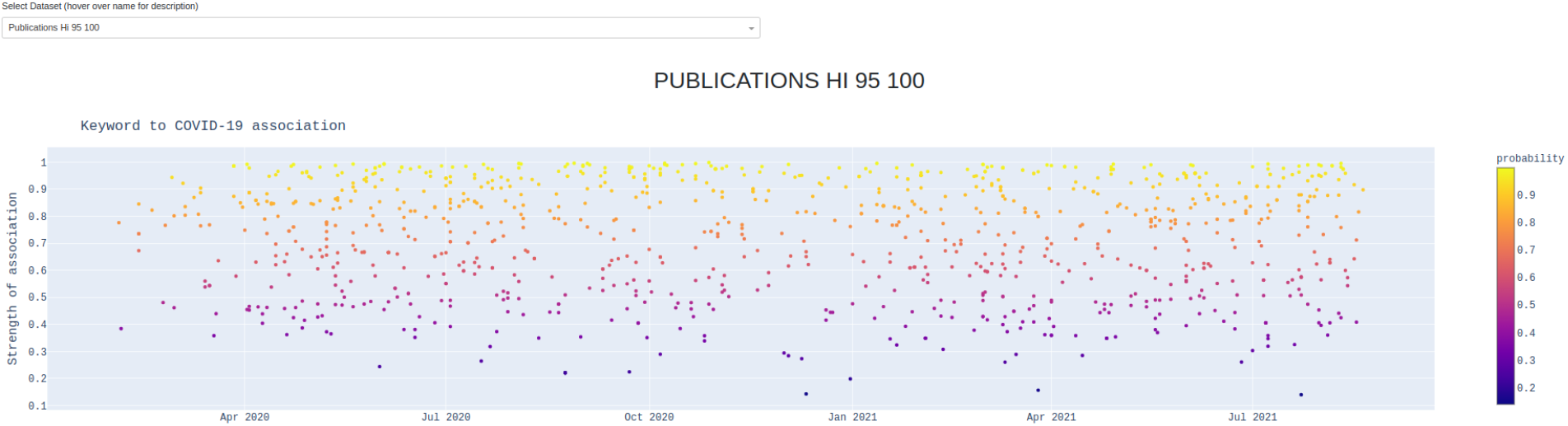
All associations between all medical categories’ keywords and COVID-19, with strength of association.

**Figure 4.**
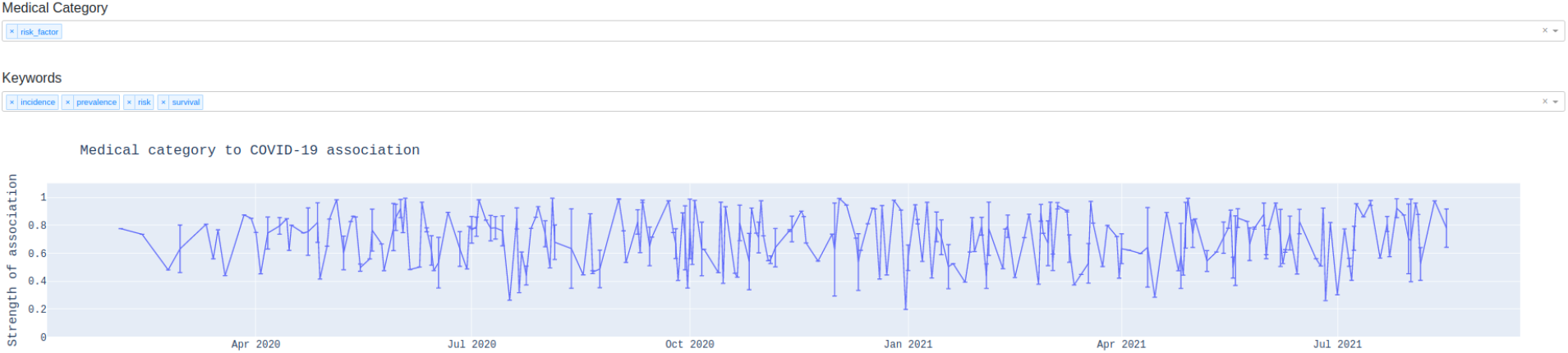
Medical category strength of association over time. Each time point depicts the averaged strength of association accompanied by 95% confidence intervals.

**Figure 5.**
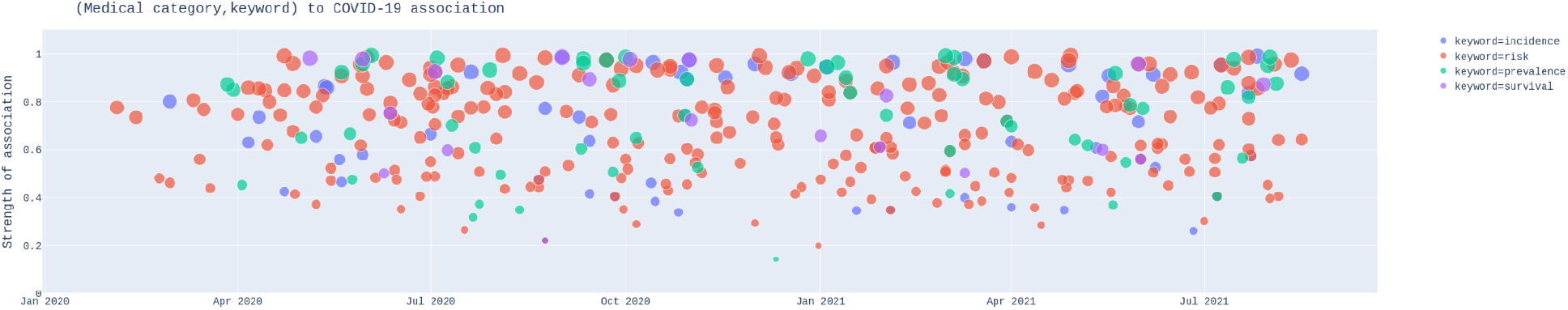
Medical category strength of association over time. Keywords are grouped into high level groups of keywords using MeSH..

The proposed framework was applied to another two areas: adolescent nutrition and lupus nephritis. The data source for both domains is pubmed articles ranging over a period of 40 years.

## 4. Conclusion

In this work, we proposed a framework for finding insights in large volumes of research datasets. We automated and combined topic modeling with a pretrained BERT encoder for insight generation, and incorporated the framework in an open source web application. Limitations of this work include the difficulty of dealing with abbreviations and contextualizing identified patterns. Future work involves the development of a user research survey for user experience analysis of the online web application.

## Acknowledgements

The authors would like to thank Dr. David Goodman for his contribution in defining the expert defined keywords for each medical category depicted on our dashboard. The authors would also like Dr. King Chung Ho for contributing in creating the pipeline of this work, the dashboard, and reviewing this article.

## Footnotes

1 http://covidinsights.herokuapp.com/

2 https://github.com/panas89/coronaBreakSuck2020/blob/master/covid/models/paperclassifier/Davids_interest.yaml

3 https://github.com/panas89/coronaBreakSuck2020

